# Converting color memory toward a spatial format to benefit behavior

**DOI:** 10.64898/2026.02.27.708515

**Authors:** Amit Rawal, Michael J. Wolff, Rosanne L. Rademaker

## Abstract

Visual working memory allows for the brief maintenance of information to serve behavioral goals. It has been shown that when the specific action required to serve a future goal is predictable, people can flexibly change a visual memory representation to incorporate an action-based one, demonstrating the goal-oriented nature of visual working memory. Can such flexibility also be observed within the visual domain, between color and space? In this eye-tracking study, participants remembered either a centrally presented color or a spatial position around fixation. Critically, when remembering a color the response wheel was either randomly rotated, or shown at a fixed rotation, on every trial. When fixed, every target color could be associated with a predictable position on the wheel during response. Do people incorporate this added spatial information in their behavior? Participants utilized color-space associations when remembering color: Response initiation happened faster when the color wheel was fixed compared to random, irrespective of whether an action could be planned or not. Next, we showed that gaze was biased towards the position of the spatial memory target during the delay, extending previous work on gaze biases. Importantly, also when remembering a color, gaze was biased towards the anticipated position of that color on the response wheel when it was fixed. Together, our results show a behavioral benefit of added spatial information for color memory, and systematic changes in gaze that reflect flexible utilization of space.

## Introduction

Imagine you’re painting a picture using colors placed on a color palette. When you have a specific color in mind, initially you have to look down at the palette to find the matching paint and then dip the brush – you rely on the appearance of the color. But as you progress with the painting, you become faster in finding the desired color, and sometimes you don’t even have to look at the palette since you can associate the color in mind with a specific location on the palette – you rely more on the spatial position. In this instance, your memory of the appearance of a color can be used interchangeably or in association with your memory of its spatial position.

Visual working memory is the ability to maintain and manipulate visual information held in mind over a short interval, without direct sensory access through the eyes. Typically, engaging your visual working memory serves a behavioral goal, such as precisely recalling a particular color, or remembering a specific position in your environment, e.g., to find a color on a palette. The passive maintenance of simple visual features (e.g., color, orientation, spatial frequency, etc.), and objects (e.g., faces, simple shapes, etc.) have traditionally received much research interest. This means that we know a lot about how recall is impacted by the number of items in memory (Luck & Vogel, 1997; Cowan, 2001), by the duration that items have to be maintained (Rademaker et al., 2018; Shin, Zou, and Ma, 2017; Magnussen & Greenlee, 1999), or by the time available for encoding (Bays et al., 2011; Sahakian et al., 2025; Giorjiani, Rawal, & Rademaker, 2026). Likewise, research on passive maintenance has revealed interactions between multiple items held in memory (Chunharas et al., 2022), between items in memory and perceptual inputs (Rademaker et al, 2015; Fukuda et al., 2022), and between items separated in time (so called ‘stimulus history’ or ‘serial bias’ effects; Fischer & Whitney, 2014). In all these cases, the behavioral requirement is straightforward – participants report the physical attributes they had to remember. While this approach has revealed much about short-term memory storage under conditions with fixed behavioral demands, in everyday life behavioral demands can change from moment to moment. Such changing demands may necessitate the mental manipulation of items in working memory (i.e., *what* is remembered), or a change in the format of the memory (i.e. *how* it is remembered) such that it can be more conveniently recalled later. Under changing behavioral contexts, might people flexibly format their memories to improve memory performance?

Recent work has explored if and how visual representations might be reformatted for storage in visual working memory. Notably, spatial features such as orientation and direction-of-motion, while distinctly represented in early visual cortex during perception, appear to be formatted in a common spatial frame once committed to memory (Kwak & Curtis, 2022; Duan & Curtis, 2024; Bae & Luck, 2018). More generally, it has been suggested that reformatting happens by way of categorization, such that the representation of an initial sensory input (e.g., blueish-green) drifts towards a category center (e.g., blue or green) once committed to memory (Panichello et al., 2019; Yan et al., 2023; Chunharas et al., 2025). Reformatting visual inputs for memory storage might happen automatically, to minimize neural resources or to stabilize information from interference. But reformatting may also serve behavioral flexibility (Lee, Kravitz, and Baker, 2013). Specifically, it is possible that information already stored in memory may undergo reformatting based on the exact demands of a specific task. Such flexible reformatting was demonstrated between the visual and motor domains in an experiment by Henderson et al. (2022). Participants performed a spatial working memory task where a small dot was briefly shown at a random position around fixation, to be recalled after a 16 second delay. During recall, a randomly rotated disk consisting of a light gray and a dark gray semi-circle was shown, and participants reported which side of the disk overlapped with the remembered position. The crucial manipulation was the presentation of a preview disk early during the delay. On half the trials, participants knew this preview disk would match the final disk, allowing them to infer which button to press later during recall. Results showed that when participants could anticipate which button to press, delay-period decoding of spatial working memory in early visual cortex decreased, while decoding of the prospective button press emerged in (pre-) motor areas. This study demonstrates that when faced with trial-to-trial differences in recall requirements, people can adaptively reformat the contents of their working memory in response to these changing task demands.

Similar visual-to-motor reformatting was shown in another recent study (Bae & Chen, 2024) using electroencephalography (EEG). The authors used a standard working memory task where first a random target color was shown at the center of the screen, and after a short delay, participants reported this color by clicking on a color wheel placed around fixation. In one condition, the color wheel was randomly rotated on every trial, as is the default in a task like this (Wilken & Ma, 2004; Zhang & Luck, 2008). In this case, only the color of the memory target is relevant for response. In a second condition, however, the color wheel was shown at the same fixed rotation (i.e., each color on the wheel occupied the same spatial position) from trial to trial. In this case, participants could use spatial information (e.g., “red is always on top of the color wheel”) in addition to color information (e.g., “the color was darkish red”) to prepare for a motor response at recall. The authors found that when the rotation of the color wheel was fixed from one trial to the next, participants’ recall accuracy was higher and their responses were faster. Importantly, while multivariate EEG patterns in both conditions contained color-specific signals when the memory target was on the screen, only when the color wheel was fixed did the authors find sustained above-chance decoding during the delay. The authors attributed this to the reformatting of color information to an action-oriented format.

Both these studies show that when faced with changes in task demands, visual working memory contents (of e.g., space or color) can be flexibly reformatted to an action-based format for prospective motor output. But could working memory contents be reformatted in more subtle ways, such as from one visual feature to another? If so, it might be the case that reformatting of information held in mind is a ubiquitous operation that can be applied flexibly to aid a wide variety of behaviors, action-oriented and otherwise. Here, we tested this via two experiments similar to the Bae & Chen (2024) study, where we asked participants to remember a color, and manipulated the rotation of the color wheel shown at recall (either random or fixed across blocks of trials). We asked whether participants would take advantage of the extra spatial information during blocks with a fixed color wheel by shifting the contents of their working memory away from pure color information, to also include spatial information. We operationalized the use of spatial information by tracking participants’ eyes, as it has been shown that eye movements can carry meaningful information about a memorized stimulus (van Ede et al., 2019; Linde-Domingo & Spitzer, 2024; Dong et al., 2025). As a baseline for spatial memory usage, we included a third condition where only a spatial position (not a color) had to be remembered and recalled. Crucially, we dissociated action-oriented from purely spatial responses by allowing participants to anticipate (Experiment 1) or not to anticipate (Experiment 2) the action required during their response with the fixed color wheel.

To preface our results, here we show that when participants could anticipate the rotation of the color wheel during recall, they were able to take advantage of the fixed mapping between color and space: Response onsets were faster (statistically significant in both experiments), and recall errors lower (statistically significant in the first experiment). Critically, when the color wheel was shown at a fixed rotation from trial to trial, participants’ eye position during the delay was biased towards the spatial position associated with the color they were remembering. The presence of this effect also in the second experiment, which did not allow for action planning, shows that participants were using spatial information during the working memory delay despite this information being uncoupled from a future plan for action. Together, these findings suggest that information in visual working memory may not only reflect the passive storage of previous sensory input (e.g., color), it may reflect visual features that are flexibly integrated when they are relevant for behavior (e.g., spatial position). This work provides further support for the idea that task demands influence the format by which working memory contents are stored (Henderson et al., 2022; Bae & Chen, 2024).

## Experiment 1

### Methods

#### Participants

39 total participants between the ages 18 and 35 were recruited for this experiment (27 female, mean age = 23 + 3.36 years old). Three participants were excluded from analyses due to >30% missing eye data (see **Eye data analysis** below). All participants had normal or corrected-to-normal vision and received monetary compensation of €12.50 per hour. Participants provided written informed-consent with approval from the Institutional Review Board at the medical department of Goethe University, Frankfurt, Germany. The experiments took place at the Ernst Strüngmann Institute (ESI) of the Max Planck Society, Frankfurt am Main, Germany.

#### Stimuli and procedures

Stimuli were generated on a Linux Red-Hat Enterprise OS (RHEL8) machine using the Psychtoolbox 3 and Eyelink toolboxes (Brainard, 1997; Kleiner, et al., 2007; Cornelissen, Peters, & Palmer, 2002) under MATLAB R2021b (The MathWorks Inc., Natick, MA, USA). Stimuli were presented on a 39.5 x 29 cm ViewSonic G225f CRT monitor (refresh rate = 85Hz, screen resolution = 1600×1200) against a uniform gray background (29.8 cd/m^2^). Participants sat 74 cm away from the screen in an otherwise dark room, head-stabilized using a chin and forehead rest. To record responses, a Sony DualShock 3 controller was used.

Eye position and pupil size data were recorded for both eyes at 1000 Hz, using an Eyelink 1000 Plus (SR Research Ltd., 2013). Prior to the experiment, a 9-point calibration was performed, followed by a validation procedure to confirm stable and accurate calibration. Saccades, blinks, and periods of fixation were detected using the default SR Research algorithm. To align continuous eye data with task events, timestamped event markers sent by the stimulus presentation computer were saved in the Eyelink data files (EDFs). Drift checks were performed before every experimental trial to ensure continued calibration accuracy, with drift deviations kept under 1 degree of visual angle (d.v.a.).

Participants performed a visual working memory task (**Figure 1**) where the to-be-remembered target stimulus, shown for 250 ms, was either the color of a centrally-presented disk (radius = 1.87°), or the spatial position of a peripherally-presented white dot (radius = 0.62°, shown at 5.61° from fixation). The target stimulus was pseudo-randomly chosen to be one of 360 evenly spaced hues spanning the HSV space at full saturation and brightness (in case of color memory), or one of 360 possible angles relative to fixation (in case of spatial memory). To reduce possible afterimages, the target stimulus was directly followed by a 250 ms centrally presented mask (radius = 8.41°), consisting of randomly colored tiles of 0.19° by 0.19° each. Participants subsequently remembered the exact color or spatial position over a delay period of 2500–3000 ms (randomly jittered). After the delay, a 0.37° wide response wheel was shown at a radius of 5.61° from fixation, and participants could move the left joystick on the game controller to indicate their response. In the case of color memory, the response wheel consisted of 360 discrete colors, while for spatial memory the wheel was dark gray. Only during color memory trials, a light gray disk appeared centrally at the onset of the recall phase (radius = 1.87°). Once a movement with the joystick was initiated, a response line (color memory, 0.71° long) or dot (spatial memory, 0.62° radius) appeared on the wheel, indicating where participants moved the joystick. In the case of color memory, the color of the central light gray disk switched to the color selected by the joystick, and tracked the selected color throughout the response period. There was no time limit imposed on the response, which allowed participants to finetune their responses as precisely as possible. Participants confirmed their final response by pressing the ‘X’ key. The next trial started after a drift-check and inter-trial-interval of 800–1000 ms. Throughout the task, participants fixated on a centrally presented fixation dot (0.19° white dot with 0.15° black dot superimposed).

**Figure 1:**
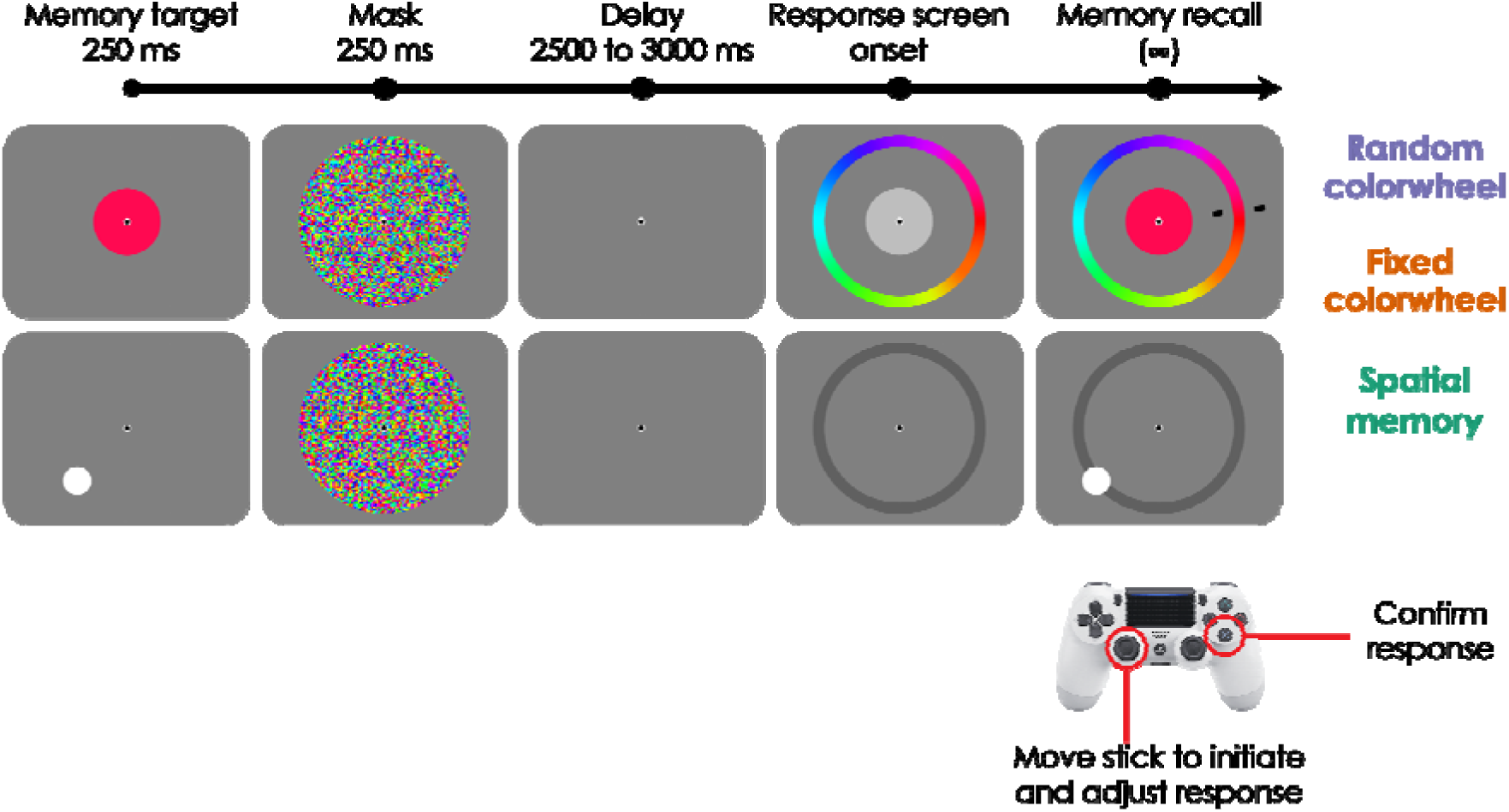
Trial sequence in Experiment 1. Each trial began with the presentation of the memory target for 250 ms, which was either a randomly chosen color (top) or spatial position (bottom), depending on the experimental block. Next, a colorful mask was shown for 250 ms, and participants kept the memory target in mind over a blank memory delay of 2500–3000 ms. A response wheel was shown around fixation, which consisted of 360 colors and a centrally presented gray disk (color memory), or was dark gray without a centrally presented disk (spatial memory). A directional movement with the controller joystick initiated the response, and a line (color memory) or dot (spatial memory) would appear on the wheel to indicate the selected response. This line/dot continued to indicate the corresponding position on the wheel as participants fine-tuned their response via the joystick in an unspeeded manner until confirmation by a keypress. In color memory blocks, the centrally presented disk would track the selected color throughout the response phase. Participants completed three different blocks of trials. In two of these blocks, they remembered a target color, and we manipulated whether the response wheel was randomly rotated across trials (the ‘random color wheel’ condition) or whether the response wheel was identical on all trials of a block (the ‘fixed color wheel’ condition). Note how in the fixed color wheel condition each target color could be associated with a specific position on the response wheel. In a third block of trials, participants remembered a spatial position. The order of the three blocks was counterbalanced across participants.

In total, there were three distinct conditions, two color memory conditions, and one spatial memory condition. These three conditions were presented in separate blocks (of 120 trials per block), and the order of blocks was counterbalanced across participants. The two color memory conditions were identical until the response phase, and only varied in one respect: The response wheel, with its 360 colors, was either randomly rotated on every trial in a block (the ‘random color wheel’ condition), or its rotation was kept fixed throughout all trials in a block (the ‘fixed color wheel’ condition). Specifically, in the ‘fixed color wheel’ condition, each of the 360 colors retained the exact same spatial position on every trial of the block (e.g., purple always appeared on top, light blue always on the left, lime green always on the bottom, etc.). For each participant a different fixed rotation was pre-selected and applied throughout, such that the color-to-space mapping in this condition was roughly uniform across all participants. Importantly, blocks of trials with a fixed color wheel allowed participants to associate every target color with a specific spatial position during recall. If visual working memory can be flexibly reformatted, incorporating such spatial information could be used to benefit behavioral performance.

### Analyses

#### Behavioral data analyses

Did the remembered visual feature (color or spatial position), or the potential use of a color-space association in the fixed color wheel condition, affect memory recall? We computed two performance metrics based on the angular responses collected from the game controller. First, we computed the ‘initial landing error’ as the absolute angular distance between the correct position of the memory target on the response wheel, and the first position selected by participants. The second metric, ‘recall error’, is the absolute angular distance between the correct position of the memory target on the response wheel, and the final position selected by participants. Average performance metrics for each participant and condition are used for statistics and plotting. Recall speed was similarly analysed by computing the ‘response onset time’, defined as the time between response wheel onset and response initiation, and ‘response adjustment time’ defined as the time between response initiation and response confirmation. Median response times for each participant and condition are used for statistics and plotting. Statistics were conducted via one-way repeated-measures ANOVAs, with Greenhouse-Geisser corrected p-values reported for cases where the assumption of sphericity was violated. Statistically significant main effects were followed by pairwise post-hoc Holm t-tests.

### Eye data analyses

#### Preprocessing

After data acquisition, the EDFs were converted to ASCII using the GUI-based *edf2asc* tool (SR Research Ltd., 2013). These data were read into MATLAB and converted to FieldTrip Toolbox format (Oostenveld et al., 2011; Urai, Braun, & Donner, 2017) for further analysis. The continuous pupil time-series data from each eye were used to adjust the onsets and offsets of Eyelink-detected blinks, which we did to account for distortions induced by partial occlusion of the pupil just before and after blinks (Hershman, Henik, & Cohen, 2018). The pupil data during these blink periods were linearly interpolated. All pupil data were then low-pass filtered at 10 Hz using a 2^nd^ order Butterworth filter in order to attenuate high-frequency noise. The continuous eye position data were kept as-is (i.e., no interpolation or filtering was applied).

Next, continuous pupil and eye position data from both eyes were segmented into ‘encoding and delay’ and ‘response’ epochs. The encoding and delay epochs consisted of data from 800 ms before the onset of the memory target until 3000 ms after it (i.e., from pre-stimulus fixation until the end of the delay period). The response epochs consisted of data from 2000 ms before the onset of the response wheel until 2000 ms after it (i.e., including part of the delay period as well as the response period). The epoched pupil data were baseline-corrected by dividing each timepoint in a given epoch by the average pupil size during the first 50 ms following that epoch’s onset (Mathôt, & Vilotijević, 2023), and were subsequently downsampled to 100 Hz for ease of computation. These processed pupil epoch data were then used to assess data quality and determine participant exclusion. Specifically, we computed the overall proportion of missing data (due to blinks) separately for the encoding and delay and response epochs of each experimental condition, and excluded participants who had >30% missing pupil data across all trials, in either epoch type of any of the three experimental conditions.

Eye position epochs were corrected for blink-induced distortions by marking blink periods (as detected on the pupil time series) as NaN. Upon visual inspection, this correction was found to be insufficient and we marked an additional 50 ms before blink onset and after blink offset. Additionally, spikes in the epoched eye position data were linearly interpolated. Specifically, spikes were detected as any sample in the 1000 Hz data which exceeded its neighboring samples by 0.7° of visual angle (i.e., an instantaneous velocity of 700° s^-1^). The resulting epoched eye position data were used in the gaze bias analysis, as described next.

#### Gaze bias analysis

To see if participants’ eye position carried spatial information about the task, we computed gaze bias throughout the entire trial (both epochs) for all three conditions. Gaze bias is defined as movement of the eyes *towards* the position of the memory target on the response wheel (the ‘target position’). Note that during color memory blocks, the target position on the response wheel can (fixed color wheel) or cannot (random color wheel) be anticipated once the target color is shown. Specifically, gaze bias was calculated by first averaging data from the two eyes, and then computing the timepoint-by-timepoint Euclidean distance (in °) between the eye position and the target position. We applied a baseline-correction to all time points by subtracting the average distance between eye- and target position in the 100 ms prior to the memory target onset. The same baseline was used for both encoding and delay and response epochs, to avoid possible gaze biases during the delay period spilling over into the response epoch. Epochs with baselines that included blinks were excluded from analysis. In the baseline-corrected distances, negative values (smaller distances) indicate an attraction *toward*, while positive values (larger distances) indicate repulsion *away* from the target position.

Because the number of trials in each condition (120) is considerably smaller than the number of degrees (360) on the response wheel, random sampling may have resulted in non-uniformity of target positions across space. To make sure that this did not impact the average gaze bias, we uniformly binned the 360° response wheel into 12 bins (of 30° each), and determined which trials (based on their target position) fell into each bin. Because not every bin contained the same number of trials, we subsampled the trials in every bin to the minimum trial number across all bins before averaging the timepoint-wise gaze bias. This ensured approximate uniformity of target positions across space. We repeated this procedure 100 times, i.e., randomly subsampling a new set of trails on every iteration. We averaged the resulting 100 gaze bias time courses to obtain a final gaze bias time course for each participant. Each participant’s average gaze bias was then temporally smoothed by computing a moving average using a 50 ms sliding window.

Statistics were conducted by means of one-sample cluster-based permutation tests (Maris & Oostenveld, 2007), performed separately for each epoch type and each experimental condition. Briefly, these tests first generate null distributions by randomly sign-flipping each participant’s gaze bias over 10000 permutations. On each permutation, this yields a shuffled *t*-value for each time point, and a shuffled sum of *t*-values indicating the largest cluster across time points (with clusters identified as contiguous timepoints during which *t*-values meet the cluster formation criterion of *p* < 0.05). To find significant clusters of gaze bias in the intact data, first the true *t*-value at each timepoint was compared to the distribution of shuffled *t*-values at that timepoint. Clusters are then identified in the intact data, after which each cluster’s sum of *t*-values is compared to the distribution of shuffled summed *t*-values. Clusters that meet the criterion of *p* < 0.05 relative to the cluster-level null distribution are considered statistically significant.

### Results

In this experiment, participants completed three blocks of a visual working memory task. In two of the blocks, participants remembered the color of a centrally presented target. Critically, while the ‘random color wheel’ condition did not allow for an association between the target color and its spatial position on the response wheel (the color wheel was rotated randomly from trial to trail), in the ‘fixed color wheel’ condition such an association *was* possible (the color wheel had the same rotation on every trial). Did participants incorporate this additional source of spatial information into their responses? To establish a baseline of spatial memory responses, a third experimental block asked participants to remember the spatial position of a peripherally presented target. Note that in this first experiment the spatial position of the target on the response wheel is directly linked to the action of recalling that target. Thus, when the target position can be anticipated prior to recall (i.e., in the spatial and fixed color wheel conditions), this enables participants to plan a movement in its direction (as in Bae & Chen, 2024). Can we verify that recall performance and reaction times are impacted by the ability to utilize such spatial information?

A spatial position held in working memory (e.g., to the left or right of fixation) can be reflected in systematic changes in participant’s eye position during the delay, namely, a ‘gaze bias’ toward the remembered position (van Ede et al., 2019). We analysed the eye position data in all 3 blocks to (1) verify the absence of systematic eye movements towards the target position on the color wheel when no association between color and space can be made, (2) examine if participants incorporate spatial information in their gaze behavior when they are able to form color-space associations, and (3) replicate previous gaze bias findings in spatial working memory tasks, but over a full range of 360° possible positions.

#### Behavioural benefits of color-space association

During recall, participants used the controller joystick to make an initial movement towards the response wheel (usually in the direction of the target) and then fine-tuned their response. We first looked at the initial landing error (**see Methods**) and found that the three working memory conditions differed significantly (F_(1.3,45.43)_ = 27.501, *p* < 0.001, η_p_^2^ = 0.440) such that initial landing error was smallest in the spatial condition compared to the fixed (*t*_(35)_ *=* 4.81, *p* < 0.001) and random (*t*_(35)_ = 7.3, *p* < 0.001) color wheel conditions (**Figure 2A**). Interestingly, the initial landing error in the fixed color wheel condition was smaller than in the random color wheel condition (*t*_(35)_ = 2.5, *p* = 0.015), implying a benefit of the additional spatial information when the color wheel was fixed. The same was true for the final recall errors (**Figure 2B**), which also differed between conditions (F_(1.70,59.37)_ = 43.17, *p* < 0.001, η_p_^2^ = 0.55). Recall errors were again smallest for the spatial condition compared to the fixed (*t*_(35)_ = 6.55, *p* < 0.001) and random (*t*_(35)_ = 8.98, *p* < 0.001) color wheel conditions, and were smaller with a fixed compared to a random color wheel (*t*_(35)_ = 2.44, *p* = 0.017).

**Figure 2:**
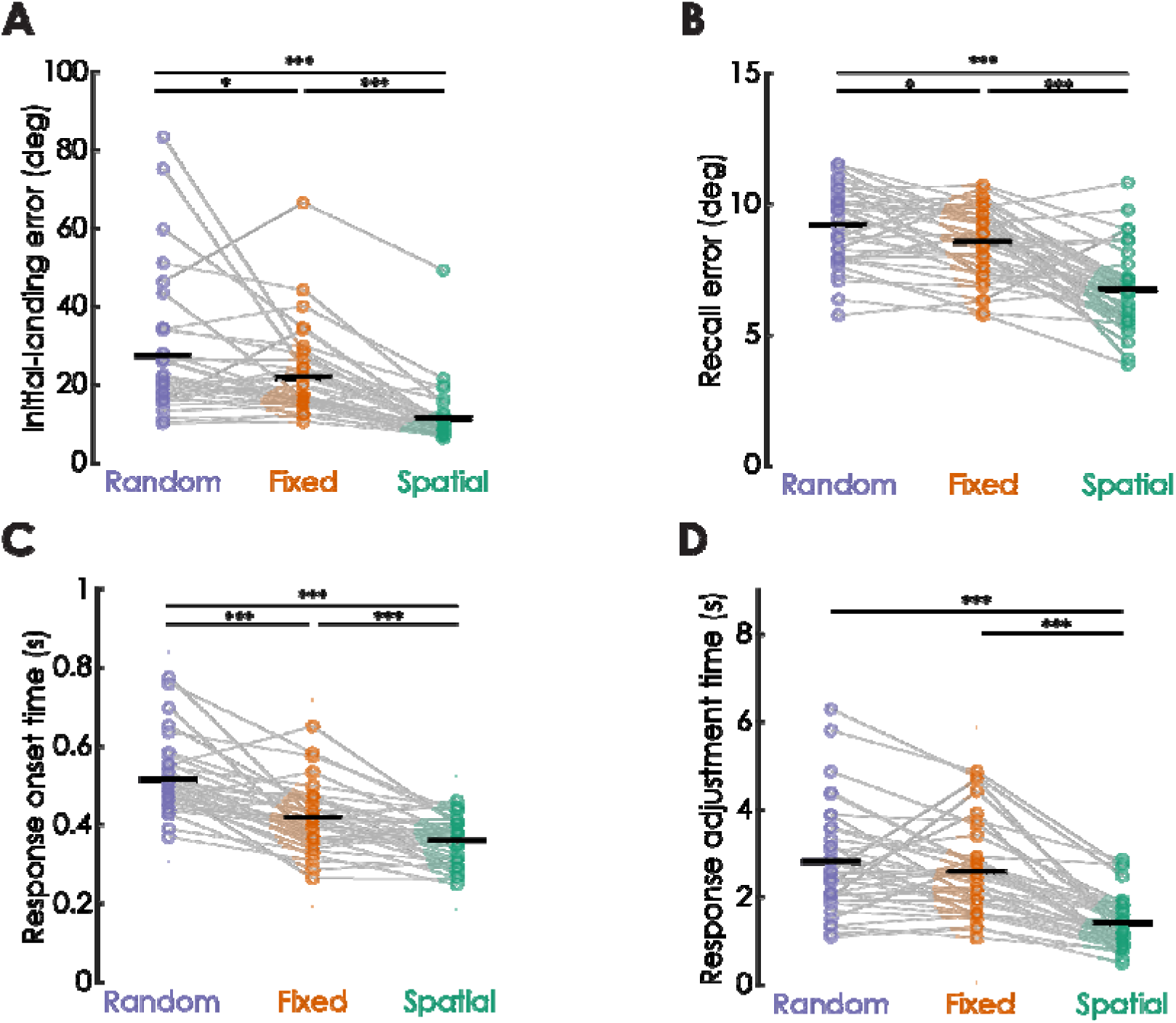
Experiment 1 task performance. Participants reported memory targets on a response wheel around fixation by first making a directional movement towards the wheel, and then fine-tuning the response via joystick rotations until confirmation via keypress. **A.** Initial landing error shows the mean absolute distance (in degrees) between the first directional movement and the target. A repeated-measures ANOVA showed that the initial landing error significantly differed across conditions (*F*_(1.3, 45.43)_ = 27.50, *p* < 0.001, η_p_^2^ = 0.44), with the lowest error in the spatial condition, and a lower error in the fixed vs. random color wheel condition. **B.** Recall error shows mean absolute distance (in degrees) between the final response confirmed on the response wheel and the target. A repeated-measures ANOVA showed that this final recall error differed significantly between conditions (*F*_(1.7, 59.37)_ = 43.17, *p* < 0.001, η_p_^2^ = 0.55), with the lowest error in the spatial condition, and a lower error in the fixed vs. random color wheel condition. **C.** Response onset time shows the median time between response wheel onset and response initiation. A repeated-measures ANOVA showed that response onset times differed significantly between conditions (*F*_(2,70)_ = 59.93, *p* < 0.001, η_p_^2^ = 0.63), with fastest onset time in the spatial condition, and faster onset time in the fixed vs. random color wheel condition. **D.** Response adjustment time shows the median time between response initiation and response confirmation, and reflects how long people fine-tuned their responses. A repeated-measures ANOVA showed significant differences in response adjustment between conditions (*F*_(2,70)_ = 46.29, *p* < 0.001, η_p_^2^ = 0.57), driven by fastest adjustments in the spatial condition. There was no difference between the color conditions, indicating people took equally long to fine-tune their color report irrespective of the color wheel being fixed or random. Post-hoc pairwise comparisons were corrected for multiple comparisons using Holm’s method, and the corrected significance levels refer to: * p < 0.05; ** p < 0.01; *** p < 0.001. Colored circles indicate individual participants, half-violins show the distribution across participants, and thick black lines show the group mean (in A and B) or median (in C and D).

Participants also had different response onset times between conditions (**Figure 2C**; F_(2,70)_ = 59.928, *p* < 0.001, η_p_^2^ = 0.631), with onset times being fastest in the spatial memory condition compared to the fixed (*t*_(35)_ = 4.05, *p* < 0.001) and random (*t*_(35)_ = 10.83, *p* < 0.001) color wheel conditions. A fixed color wheel resulted in faster response onsets compared to a random color wheel (*t*_(35)_ = 6.78, *p* < 0.001), again implying a benefit when anticipating the spatial position of a given target color on the response wheel. This pattern of results mostly repeated when looking at the time participants took to adjust and fine-tune their responses, which is defined as the time between response onset and response confirmation (**Figure 2D**). Response adjustment times differed between conditions (F_(2,70)_ = 46.294, *p* < 0.001, η_p_^2^ = 0.569), and were again fastest in the spatial condition (compared to the fixed wheel: *t*_(35)_ = 7.48, *p* < 0.001; and random wheel: *t*_(35)_ = 8.98, *p* < 0.001). However, the difference in adjustment times between the two color conditions did not reach significance (*t*_(35)_ = 1.5, *p* = 0.137), suggesting that while participants are faster to *initiate* their responses when they can anticipate the spatial position of the remembered color, they tend to spend a similar amount of time *fine-tuning* their final color choice irrespective of whether the color wheel is random or fixed.

Taken together, we show that memory recall is best (in terms of both accuracy and speed) when people are remembering a spatial position, compared to a color. However, when color can be recalled on a color wheel that is fixed from one trial to the next – allowing people to form color-space associations – performance is still better than when the color wheel is randomly rotated from one trial to the next. When the color wheel is random, participants cannot anticipate the position of the target color on the color wheel, and need to rely purely on color memory to locate the target during recall. When the color wheel is fixed, the approximate position of the target color on the color wheel can be known in advance, and our results imply that participants use this added spatial information to their advantage. Moreover, the fact that participants tended to be both faster and more accurate when they could take advantage of spatial information means there is no speed-accuracy trade off. Rather, participants do better all-around when they can increasingly rely on space. That said, this first experiment cannot disambiguate between two possible strategies people might have employed to take advantage of the additional spatial information: Participants may have relied on spatial information, on the ability to prepare an action, or both. To understand if these results are just another instance of visual working memory contents being cast into an action-based format (Henderson et al., 2022; Bae & Chen, 2024), or if spatial information alone is sufficient to observe these improvements in behavior, we would need to uncouple spatial information from action plan (see Experiment 2).

#### Gaze bias as an index of color-space association

To test whether the ability to rely on spatial information is also reflected in people’s ocular behavior, we used the eye position data recorded during the task to see whether there were systematic changes in eye position toward the target position on the response wheel. Subtle shifts in eye position can index a spatial position selected for working memory (van Ede et al., 2019; van Ede, Board, & Nobre, 2020; Draschkow, Nobre, & van Ede, 2022). We extend these findings to a situation in which only a single spatial position is relevant, and can be remembered at any angle around fixation. Moreover, a gaze bias may not be unique to memory targets at different spatial positions, but could also emerge in case of a color-space association. We reasoned that associating a memorized color with a specific position on the fixed color wheel could also result in an attractive gaze bias towards this position during the delay. Of course, in the absence of a color-space association (random color wheel), it should be impossible to observe a gaze bias toward the future position of the target color on the wheel. We looked at gaze bias during memory encoding and the delay (−800 to 3000 ms relative to the target onset) and when participants gave their response (−2000 to 2000 ms relative to the response wheel onset). Note that gaze bias is calculated as the Euclidean distance (in d.v.a.) between the eye position and target position on the response wheel, which means that a negative gaze bias indicates an attraction *towards* the target position (see **Methods** for more details).

During the encoding and delay epoch (**Figure 3A**) we observed an attractive gaze bias towards the target position in the spatial condition (cluster *p* < 0.001), which became apparent soon after the memory target appeared on the screen, and persisted throughout the delay. Notably, an attractive gaze bias was also observed halfway through the delay in the fixed color wheel condition (cluster *p* = 0.0374), but not the random color wheel condition (no statistically significant clusters). This shows that even during a color working memory task, and already before the onset of the color wheel, participants’ gaze drifts in the anticipated direction of the memorized color, making apparent the color-space association. During the response epoch (**Figure 3B**), when participants were initiating and fine-tuning their memory recall, we find a gaze bias for all conditions (all cluster *p’*s < 0.001) in the form of an attraction towards the on-screen position of the memory target on the response wheel. For both color conditions, there is a sharp increase in the magnitude of attraction (though still well within the foveal range) which peaks around 0.35 s after the onset of the response screen. This places the peak gaze bias shortly before the response onset time in both color conditions (fixed wheel: 0.42 ± 0.08 s; random wheel: 0.52 ± 0.09 s). For the spatial memory condition, the gaze bias during the response increases more gradually, and peaks later (around 1 s after the onset of the response screen).

**Figure 3:**
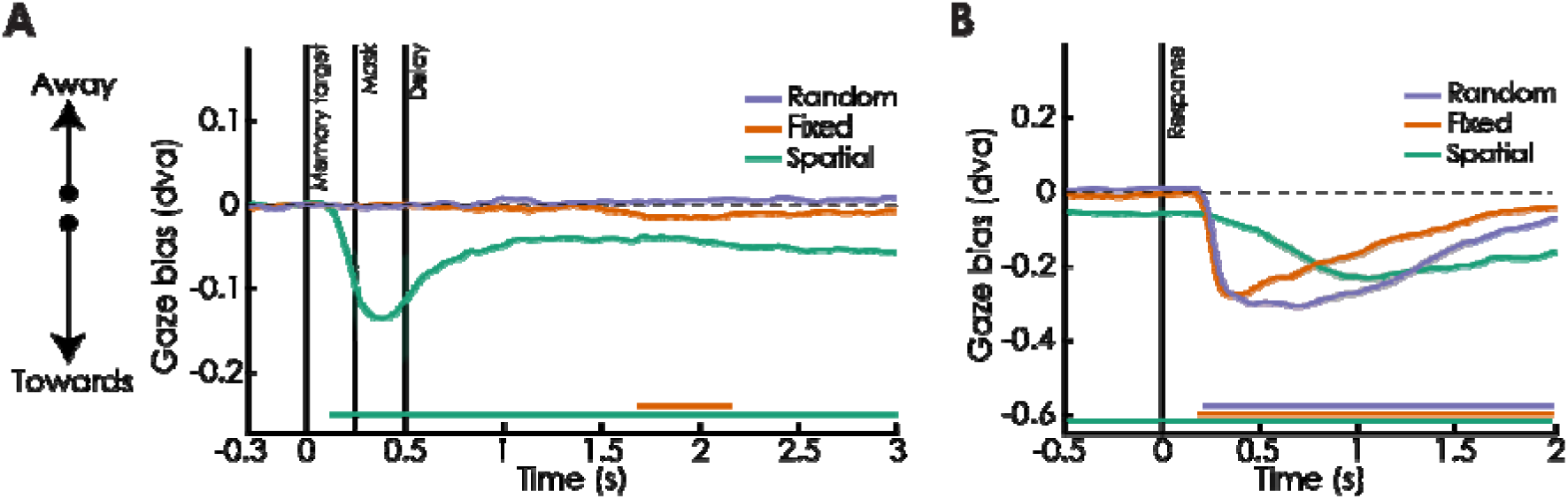
Experiment 1 gaze bias toward the target position. Gaze bias (Euclidean distance between the eye and target position, in d.v.a) is plotted over time for all three working memory conditions. Note that a negative gaze bias indicates a smaller distance (i.e., attraction) to the target position on the response wheel. **A.** During the encoding and delay epoch, gaze was biased toward the target position for the spatial condition soon after the target appeared on the screen and throughout the delay (cluster *p* < 0.001). In the fixed color wheel condition, gaze was also biased toward the target position during a portion of the delay (cluster *p* = 0.0374). These gaze biases reveal a spatial signature of a behaviorally relevant position held in mind. **B.** During the response epoch, participants’ gaze was biased toward the on-screen target position for all three working memory conditions (all cluster *p* values < 0.001). Colored lines indicate condition group average and shaded areas show bootstrapped 95% confidence intervals. Colored horizontal bars depict significant clusters in time for the respective conditions (α = 0.05), with significant clusters identified via one-sample cluster-based permutation tests (two-tailed). The dashed horizontal line marks zero (no gaze bias). Vertical lines and respective labels indicate different trial events.

## Experiment 2

In Experiment 1, we found that when people can rely on spatial information during a working memory task, their recall performance is better, response times are faster, and their gaze is biased towards the relevant spatial position. These effects are strongest when spatial position is the only feature relevant for the task (spatial condition), but they are also apparent when participants remember a specific color that they are able to associate with a specific position in space (fixed color wheel condition). However, one limitation of Experiment 1 is that the influence of spatial position cannot be dissociated from potential action planning. This is because responses in Experiment 1 were made by moving a controller joystick in the direction of the memory target. This meant that the spatial direction of a target always had a one-to-one correspondence to the direction of the motor response during recall, allowing the use of a non-visual strategy (i.e., motor preparation). In the absence of action-based strategies, can the presence of purely spatial information during a working memory task also be used to improve recall performance and bias gaze? In this second experiment we uncouple spatial information from the motor output during recall, and remove participants’ ability to generate a motor plan toward the target position. Specifically, we used a rotating dial as the response device, which removed the directional aspect of the initial response towards the target position, and broke the correspondence between motor output and selected position on the response wheel.

### Methods

#### Participants

39 total participants between the ages 20 and 40 were recruited for this experiment (22 female, mean age = 27 ± 4.31 years old). Two participants did not complete the experimental procedures; one due to drowsiness and another due to an inability to maintain fixation. Another participant was excluded from analyses due to >30% missing eye data. The 36 remaining participants were included for analyses. All participants had normal or corrected-to-normal vision and were compensated for their time (€12.50 per hour). Participants provided written informed-consent with approval from the Institutional Review Board at the medical department of Goethe University, Frankfurt, Germany. The experiments took place at the Ernst Strüngmann Institute (ESI) of the Max Planck Society, Frankfurt am Main, Germany.

#### Stimuli and procedures

The experimental stimuli and procedures were identical in all respects to the first experiment, with the following exceptions: Instead of a game controller as the input device used to record responses, here we used a custom-built dial with 24 discrete serial outputs in one rotation cycle. We added software acceleration to allow fine movements of 1 deg precision when moved slowly, and larger movements when moved quickly. Critically, as soon as the response screen appeared, an unpredictable initial dial position was already assigned to the response wheel (**Figure 4**). Across all 120 trials in a given condition, the offset between the initial dial position and the target position was controlled (i.e., the offset was 3°–180° in 3° steps, in either clockwise or counterclockwise direction). This ensured that the total offset between target and dial was the same across all three conditions, and could not impact response times. In the two color conditions, the color corresponding to this assigned position on the wheel was displayed at fixation. Only rotational movements on the dial could be used during the response, with the position of the fingers rotating the dial uncoupled from the position indicated on the response wheel. To further reduce the correspondence between motor output and position on the wheel, software acceleration to the dial output made it difficult to predict the exact angular displacement on the response wheel when making dial rotations at different speeds. Once participants were done fine-tuning their response, they confirmed their response by pressing the spacebar key on a standard keyboard. Note that using this procedure allowed us to measure recall error, response onset time, and response adjustment time (as in Experiment 1). However, we were no longer able to determine an initial landing position, as the initial position on the response wheel was already randomly assigned.

**Figure 4:**
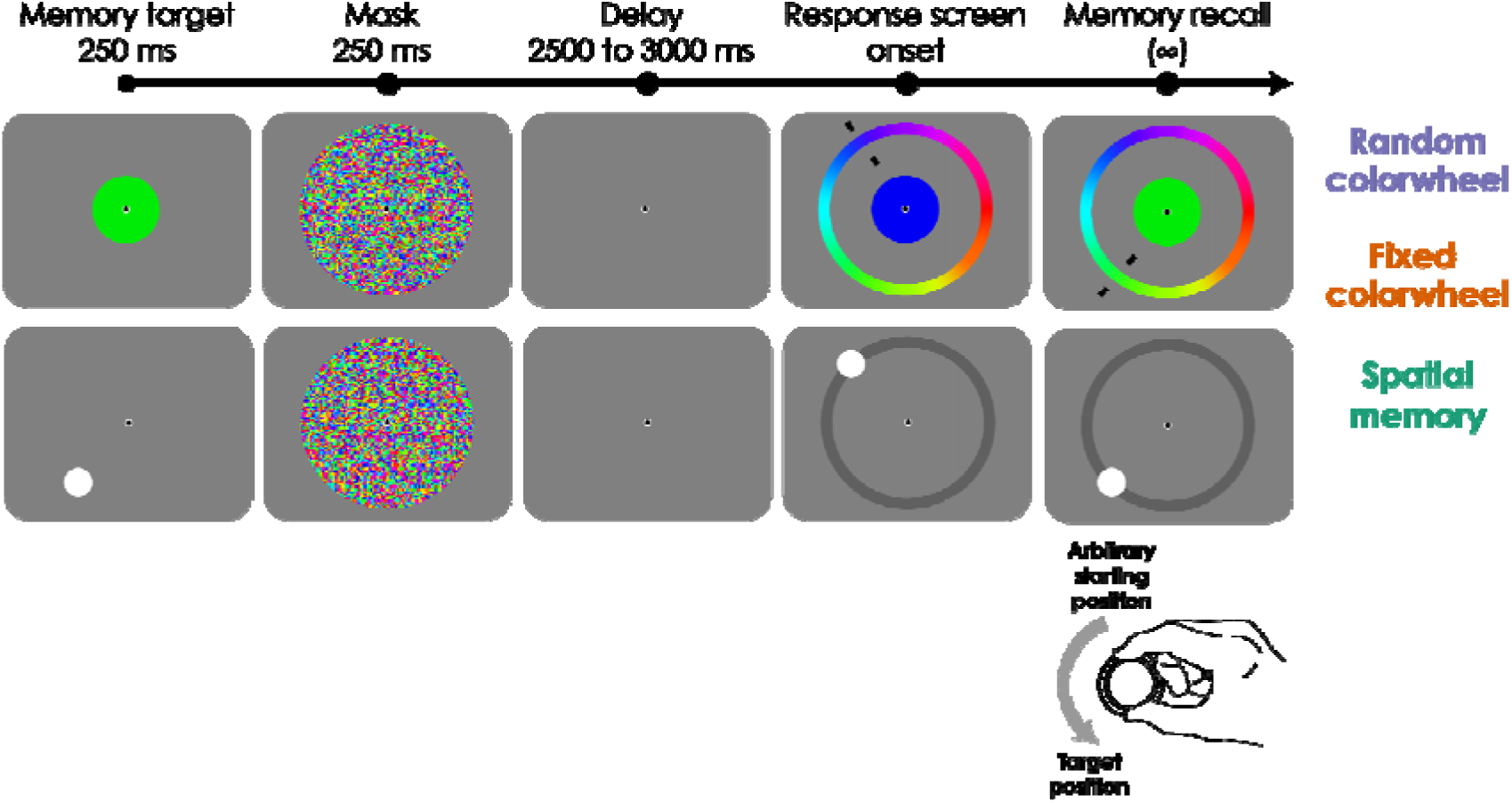
Trial sequence in Experiment 2. Each trial began with the presentation of the memory target for 250 ms, which was either a randomly chosen color (top) or a spatial position (bottom), depending on the experimental block. Next, a colorful mask was shown for 250 ms, and participants kept the memory target in mind over a blank memory delay of 2500-3000 ms. A response wheel was shown around fixation, which consisted of 360 colors and a centrally presented disk (color memory), or was dark gray without a centrally presented disk (spatial memory). At the onset of the response, a line (color memory) or a dot (spatial memory) indicated an unpredictable initial position on the response wheel. In color memory blocks, a corresponding initial color was centrally presented. Participants used a response dial to move this line/dot from its initial position to the target position in an unspeeded manner until confirmation by a keypress. The line/dot tracked the dial movements by continuously indicating the selected position on the response wheel (in color memory blocks, the centrally presented disk also tracked the selected color). Participants completed three different blocks of trials. In two of these blocks, they remembered a target color, and we manipulated whether the response wheel was randomly rotated across all trials of a block (the random color wheel condition) or whether it kept the same rotation across trials (the fixed color wheel condition). Note how in the fixed color wheel condition each target color could be associated with a specific position on the response wheel. In a third block of trials, participants remembered a spatial position. The order of the three blocks was counterbalanced across participants.

### Analyses

#### Behavioural data analyses

The behavioural data analyses in this experiment were identical to the first experiment, with the exception that the ‘initial landing error’ was not applicable here. This is because an initial position was already assigned on the response wheel at response onset, and participants only needed to rotate the response dial to match the memory target.

#### Eye data analyses

All eye data analyses were identical to the first experiment.

### Results

In the first experiment, memory recall involved making a directional movement to a position on the response wheel. In conditions where spatial information could already be anticipated during the delay (spatial and fixed color wheel) participants could have prepared an action, making it possible that some of the benefits we observed in Experiment 1 were less about space, and more about action planning. This means that also the color-space association uncovered with the fixed color wheel may really have been a color-action association. In this experiment, we remove the possibility to prepare an action, and examine if spatial information alone is sufficient to confer the behavioural benefits and gaze bias observed in the first experiment.

#### Behavioural benefits of color-space association

First, we looked at response accuracy (**Figure 5A**) by analyzing participants’ recall errors, which differed between the three working memory conditions (F_(1.54,53.86)_ = 86.473, *p* < 0.001, η_p_^2^ = 0.712). As in Experiment 1, recall errors were lowest in the spatial memory condition compared to the fixed (*t*_(35)_ = 10.9, *p* < 0.001) and random (*t*_(35)_ = 11.82, *p* < 0.001) color wheel conditions. Unlike in Experiment 1, there was no significant difference between the random- and fixed color wheel condition (*t*_(35)_ = 0.93, *p* = 0.358). This implies that a color-space association no longer benefits recall performance when it cannot be combined with an action plan. Alternatively, performance difference between the two color conditions might be difficult to detect because these differences are small (0.68° and 0.31° in Experiments 1 and 2, respectively), and performance in the second experiment was noisier overall (recall errors of 8.16° and 9.56° in Experiments 1 and 2, respectively). Such factors could have contributed to a null effect in Experiment 2, despite ⅔ of participants performing better in the fixed compared to the random color wheel condition in *both* experiments.

**Figure 5:**
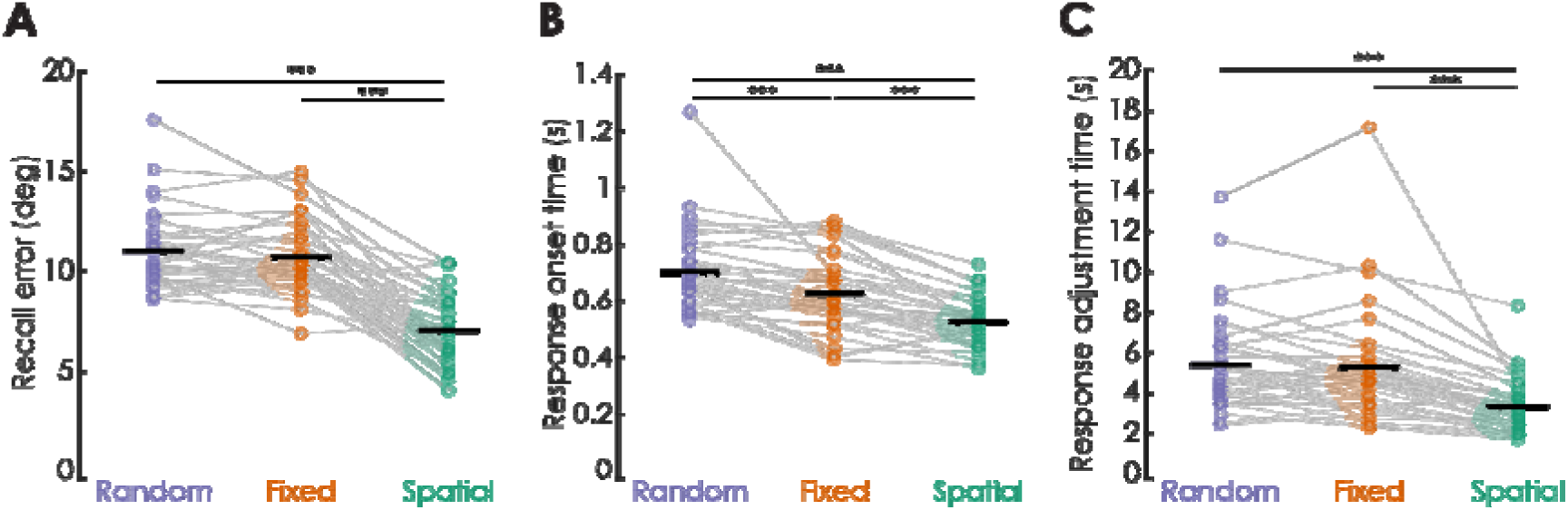
Experiment 2 task performance. Participants reported memory targets on a response wheel around fixation by rotating a dial until confirmation via keypress. **A.** Recall error shows mean absolute distance (in degrees) between the target and the final response confirmed on the response wheel. A repeated-measures ANOVA showed that recall error differed significantly between conditions (F_(1.54,53.87)_ = 86.473, *p* < 0.001, η_p_^2^ = 0.712), with the lowest error in the spatial condition, and no significant difference between the color conditions. **B.** Response onset time shows the median time between the response wheel onset and response initiation. A repeated-measures ANOVA showed that response onset times differed significantly between conditions (F_(2,70)_ = 48.409, *p* < 0.001, η_p_^2^ = 0.58), with fastest onset time in the spatial condition, and faster onset time in the fixed vs. random color wheel condition. **C.** Response adjustment time shows the median time between response initiation and response confirmation, and reflects how long people fine-tuned their responses. A repeated-measures ANOVA showed significant differences in response adjustment times between conditions (F_(1.6,55.1)_ = 32.775, *p* < 0.001, η_p_^2^ = 0.484) that were driven by fastest adjustments in the spatial condition. There was no difference between the color conditions, indicating people took equally long to fine-tune their color report irrespective of the color wheel being fixed or random. Post-hoc pairwise comparisons were corrected for multiple comparisons using Holm’s method, and the corrected significance levels refer to: * p < 0.05; ** p < 0.01; *** p < 0.001. Colored circles indicate individual participants, half-violins show the distribution across participants, and thick black lines show the group mean (in A) or median (in B and C).

Next, we looked at response times. The time that participants took to initiate their response (**Figure 5B**) differed between the three working memory conditions (F_(2,70)_ = 48.409, *p* < 0.001, η_p_^2^ = 0.58). Similar to the first experiment, response onset times were fastest in the spatial condition compared to the fixed color wheel (*t*_(35)_ = 5.55, *p* < 0.001) and the random color wheel (*t*_(35)_ = 9.8, *p* < 0.001). Importantly, here we also found quicker response onsets with the fixed color wheel than with the random color wheel (*t*_(35)_ = 4.26, *p* < 0.001). The time participants took to fine-tune their responses (**Figure 5C**) also differed between memory conditions (F_(1.57,55.09)_ = 32.775, *p* < 0.001, η_p_^2^ = 0.484). Similar to the first experiment, response adjustments were fastest in the spatial memory condition compared to the fixed (*t*_(35)_ = 6.8, *p* < 0.001) and random (*t*_(35)_ = 7.21, *p* < 0.001) color wheel conditions. Again similar to the first experiment, adjustment times in the random and fixed color wheel conditions did not differ from one another (*t*_(35)_ = 0.42, *p* = 0.677), implying that while a color-space association speeds up response initiation, it does not impact the time people take to fine-tune and adjust their responses during color recall.

In summary, we find that even in the absence of action preparation, participants used spatial information to their advantage. This was most pronounced when looking at the time people took to initiate their response, which was not only faster when spatial position was the only feature of interest (spatial memory condition), but was also faster when people could form color-space associations (fixed color wheel condition) compared to when they could not (random color wheel condition).

#### Gaze bias as an index of color-space association

Finally, we investigated whether the attractive gaze biases observed in the delay periods of the spatial and fixed color wheel conditions of the first experiment were still present when action preparation was no longer possible. Overall, we found the results in this second experiment to be comparable to those of the first experiment. Specifically, in the encoding and delay epoch (**Figure 6A**), we again found a sustained attractive gaze bias towards the spatial position of the target in the spatial condition (cluster *p* < 0.001), which emerged soon after the memory target appeared on the screen, and lasted throughout the delay. As in the first experiment, we also found an attractive gaze bias toward the spatial position associated with the remembered color in the fixed color wheel condition (cluster *p* = 0.0036), which became apparent later during the delay. No significant clusters were found in the random color wheel condition, confirming that our analysis does not lead to spurious results. In the response epoch, where the memory target was now visible on the response wheel (**Figure 6B**), we again found significant clusters of gaze bias in all three conditions (all cluster *p’*s < 0.001). As in the first experiment, the eyes were sharply drawn to the target color position on the wheel around 0.4 s after response screen onset on both color memory conditions, which preceded the time at which participants initiated their response (around 0.65 s). Gaze bias toward the target position in the spatial condition was more gradual.

**Figure 6:**
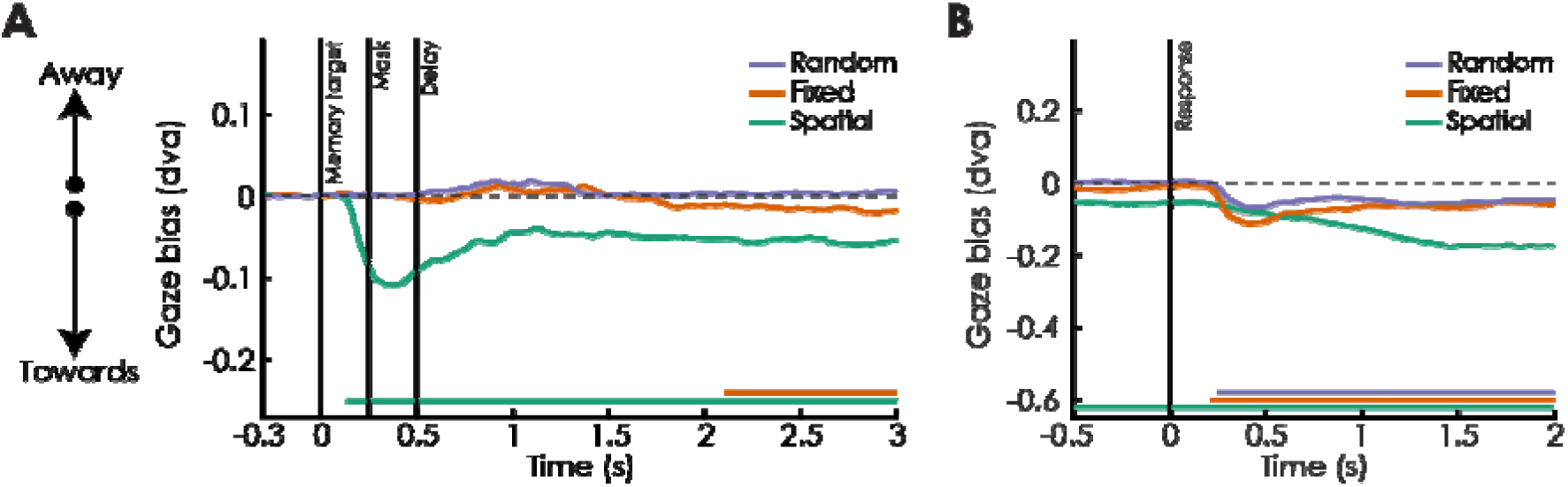
Experiment 2 gaze bias toward the target position. Gaze bias (Euclidean distance between the eye and target position, in d.v.a) is plotted over time for all three working memory conditions. A negative gaze bias indicates a smaller distance (i.e., attraction) to the target position on the response wheel. **A.** During the encoding and delay epoch, gaze was biased toward the target position for the spatial memory condition soon after the target appeared on the screen and throughout the delay (cluster *p* < 0.001). In the fixed color wheel condition, gaze was also biased toward the position of the target color on the response wheel, later during the delay (cluster *p* = 0.0036). These biases imply that the eyes carry information about the spatial position of a target held in mind, even when that spatial information was not an original feature of the target during encoding (as in the fixed color wheel condition). **B.** During the response epoch, gaze was biased toward the on-screen target position for all three working memory conditions (all cluster *p* values < 0.001). Colored lines indicate condition group average and shaded areas show bootstrapped 95% confidence intervals. Colored horizontal bars depict significant clusters in time for the respective conditions (α = 0.05), with significant clusters identified via one-sample cluster-based permutation tests (two-tailed). The dashed horizontal line marks zero (no gaze bias). Vertical lines and respective labels indicate different trial events.

## Discussion

In this study, we explored whether information stored in visual working memory can undergo a transformation from one visual feature (color) to another (space). Across two experiments, we manipulated whether participants could associate memorized colors with specific spatial locations occupied by those colors during response. This association was achieved by either keeping the response wheel at a fixed rotation across all trials (fixed color wheel condition) or by presenting it at a random rotation on every trial (random color wheel condition). As a baseline for spatial memory, participants remembered a spatial target which was then recalled on a response wheel. Across two experiments we found that memory recall was initiated faster when colors could be associated with a spatial position (fixed color wheel condition), as opposed to when they could not (random color wheel condition). This indicated the use of the additional spatial information to improve behavior. Memory recall in the spatial condition was faster and more accurate than in both color conditions.

Importantly, the spatial content of people’s working memories was also reflected in their gaze: Not only when the memorandum was a spatial target (spatial memory condition), but also when it was a color associated with a spatial position (fixed color wheel condition), we observed a gaze bias toward the position of the target on the upcoming response wheel. Together, these results show that participants can shift the contents of their working memory – relying on spatial information in addition to color information.

In the first experiment, participants could take advantage of the spatial information inherent to the memory target in the spatial memory and fixed color wheel conditions (but not in the random color wheel condition) either by using its anticipated spatial position on the response wheel, or by preparing an action in its direction. Action planning was possible due to the one-to-one mapping between movement of the joystick and target position on the response wheel. Irrespective of whether participants used visual or action-related strategies, evidence that spatial information is more conveniently recalled comes from the fact that responses in the spatial memory condition were better than in both color conditions across all performance metrics. Evidence that participants incorporated spatial information when remembering a color that could be associated with a specific spatial position, comes from their initial movement towards the color wheel. This initial movement took off more quickly and landed closer to the target color in the fixed color wheel compared to the random color wheel condition. While participants used the same amount of time to fine-tune their responses in both color conditions, the spatial benefit of a fixed color wheel remained apparent from their final response, which was also closer to the target in the fixed color wheel condition. These findings replicate the faster and more accurate responses found in an almost identical task (Bae & Chen, 2024), though this previous work did not analyze initial and final responses separately. Together, these results imply that additional spatial information in the fixed color wheel condition may reduce the overhead of locating the target color during response, allowing participants to make faster and more precise initial movements.

As our primary aim was to study the transformation between two *visual* features, color and space, we excluded the use of action-related strategies in the second experiment by breaking the one-to-one mapping between a directed action and the memory target’s position on the color wheel. Thus, the only additional information provided by the fixed color wheel was the association between colors and spatial positions. Here we uncover that participants were able to incorporate spatial information in their memory of a target color, even when no action could be planned in the delay. This advantage of added spatial information was reflected in faster response onset times with the fixed compared to the random color wheel. As in Experiment 1, exactly ⅔ of participants had a lower recall error in the fixed vs. random wheel condition, but this difference did not reach statistical significance in Experiment 2. This lack of a difference in recall error may have been the result of participants’ inability to prepare an action. Alternatively, it is also possible that this difference did not reach significance due to the smaller effect size (0.68° and 0.31° in Experiments 1 and 2, respectively) and overall noisier responses in the second experiment (recall errors of 8.16° and 9.56° in Experiments 1 and 2, respectively). As in Experiment 1, the time participants took to fine-tune their responses did not differ between the two color memory conditions. Finally, also in this experiment we saw better and faster responses in the spatial memory condition than in both color conditions.

In addition to examining participants’ memory recall, we also looked for signatures of the utilization of spatial information in their ocular behavior. Previous work has shown that even when people are asked to maintain fixation, subtle eye movements can carry meaningful information about a memorized stimulus. For example, the memorized orientation at which an object was presented can be decoded from eye positions (Linde-Domingo & Spitzer, 2024), and the spatial position of a mentally selected item can be inferred from a gaze bias towards that item (van Ede et al., 2019, van Ede, Board, & Nobre, 2020; Draschkow, Nobre, & van Ede, 2022; Liu et al., 2022). However, studies that have looked at gaze biases typically show 2-4 items at fixed spatial positions, and subsequently require participants to select one of these items from memory for later report. In the spatial memory condition from both our experiments, we reveal that gaze is also biased in the absence of any attentional competition (i.e., gaze bias toward a *single* target), and can be observed over a 360° space (i.e., the target could be presented at *any* angle around fixation). This gaze bias emerged soon after the spatial memory target (a dot) was shown, and persisted throughout the delay. The gaze bias we observed manifests as a small magnitude change in eye position, likely indicating covert spatial attention. Note that this gaze bias is relatively small when compared to what has been found in previous work. This might be because eye movements are preferentially directed along the horizontal plane (Gilchrist & Harvey, 2007; Foulsham, Kingstone, and Underwood, 2008), which happens to be where memory targets are placed in much previous work on gaze biases. Instead, the gaze bias in our paradigm relies on subtle eye movements towards a target in any direction from fixation, which may dilute the bigger effects that can be observed when targets are only placed along the horizontal meridian.

In addition to replicating that gaze is biased towards the position of a previously shown spatial memory target, both our experiments reveal that gaze can also be biased during the delay of a color memory task. Specifically, when a specific color could be associated with a specific spatial position (relevant only during the response phase), people’s gaze was biased toward this *anticipated* position. In other words, eye movements during the delay indexed the anticipated position of a memorized color (in the fixed color wheel condition). When the position of a target color at response was impossible to predict (in the random color wheel condition), no gaze bias was observed. We hypothesize that these biases reveal a transformation of the working memory contents from being purely color-based to incorporating the added (visuo-) spatial information relevant for behavior during recall.

The fact that the gaze bias in the fixed color wheel condition emerges relatively late in the delay, suggests an anticipatory orientation towards the spatial position that the memorized color will soon occupy on the color wheel. This is in contrast to the spatial memory condition where we observed a strong orienting response that began soon after the onset of the memory target. The gaze bias observed in the presence of a color-space association is notably different from most previous work in another way as well: In the fixed color wheel condition, the gaze bias does not indicate the spatial position of a peripherally presented target (in the past), but instead reflects something about a centrally presented color (in the future). Moreover, the link between color and spatial position was implicit and not part of the instructions.

Our findings complement those from a recent EEG study by Bae and Chen (2024), who used a highly similar task to study whether a non-spatial feature (color) can be reformatted to an action-oriented format. The authors were able to decode the memory target from multivariate EEG patterns during the memory delay only when the color wheel was shown at a fixed rotation from trial to trial. When the wheel was random instead, color decoding quickly returned to chance after the target was no longer on the screen. The ability to decode during the delay when the wheel was fixed was attributed to the target color being reformatted from a color representation to a more efficient action-oriented format, as participants knew in advance what movement to make during report. Thus, this experiment is most comparable to the first experiment of the present study, as it also allowed an action-based strategy to supplement color memory. As in our first experiment, such a design cannot directly disentangle to what extent spatial signals are due to action preparation, spatial attention, or a combination of both. Bae & Chen (2024) attempted to address this by looking at EEG alpha band (8–12 Hz) signals, which are thought to reflect allocation of spatial attention. When the color wheel was fixed, they could decode the memory target only during a brief window in the delay. While this finding implies the involvement of covert attention, the authors argued that color was primarily reformatted into an action-oriented format. After all, the alpha band decoding was too insubstantial to explain the sustained delay period decoding from their previous analysis (which had used EEG signals from which the alpha band activity was explicitly filtered out). However, in Experiment 2 of our current study, we removed the possibility to plan an action ahead of the response, and nevertheless found gaze biases comparable to those of Experiment 1. This suggests that gaze is biased as a result of a covert orientation towards parts of space associated with a target color, relevant during the response, but is not necessarily related to a planned action.

Unlike in our two experiments, Bae & Chen (2024) did not report systematic involvement of eye movements. Rather, they checked that eye movements did not contribute to their EEG decoding results by removing trials with eye movements (>1.5 d.v.a.), and by filtering out eye movement related signals (with independent component analysis). While this shows that these EEG results also hold without taking eye movements into account, results from the present study show that eye movements are nevertheless a valuable source of spatial information in this kind of task.

Results from the present study add to a growing body of literature showing that the contents of working memory can be flexibly reformatted depending on task demands (for review, see Adam et al., 2025; Kiyonaga & Serences, 2025). This ability to reformat implies that behavioral relevance may be a primary determinant of working memory representations (Myers, Stokes, and Nobre, 2017; Nobre & Stokes, 2019; Boettcher et al., 2021; Trentin et al. 2024), more so than perceptual properties of target stimuli. Reformatting may take the form of cross-modal (visual to action) changes in memory format (Henderson et al., 2022; Bae & Chen, 2024), or it may involve reformatting one visual feature to another, as we show here, suggesting that reformatting of information held in mind may well be a ubiquitous operation that can be applied flexibly to aid behavior.

## Acknowledgements

We’d like to thank Lea Kërçiku, Johanna Diehl, and Josephine Hahn for their help in collecting the data in this paper. AR used ChatGPT (OpenAI) for critique on the first draft (not text generation) and for code troubleshooting. MJW and RR used no generative AI tools.

